# A noncanonical repressor function of JUN restrains YAP activity and suppresses YAP-dependent liver cancer growth

**DOI:** 10.1101/2023.08.20.554005

**Authors:** Yuliya Kurlishchuk, Anita Cindric Vranesic, Marco Jessen, Alexandra Kipping, KyungMok Kim, Paul Cramer, Björn von Eyss

**Affiliations:** Transcriptional Control of Tissue Homeostasis Lab, Leibniz Institute on Aging, Fritz Lipmann Institute e.V., Beutenbergstr. 11, 07745 Jena, Germany; Leibniz Institute on Aging, Fritz Lipmann Institute e.V., Beutenbergstr. 11, 07745 Jena, Germany

**Author notes:** These authors contributed equally to the work (Yuliya Kurlishchuk, Anita Cindric Vranesic, Marco Jessen).

**Keywords:** YAP, TAZ, Hippo, AP-1, JUN, liver cancer, hepatocellular carcinoma

## Abstract

Yes-associated protein (YAP) and its homologue, transcriptional coactivator with PDZ-binding motif (TAZ), are the main transcriptional downstream effector of the Hippo pathway. Decreased Hippo pathway activity leads to nuclear translocation of YAP/TAZ where they interact with TEAD transcription factors to induce target gene expression. Unrestrained YAP/TAZ activity can lead to excessive growth and tumor formation in a short time, underscoring the evolutionary need for tight control of these two transcriptional coactivators. The AP-1 complex binds together with YAP/TAZ to many common sites and they form a positive feed-forward to induce gene expression.

Here, we report that the AP-1 component c-JUN acts as specific repressor of YAP/TAZ at joint target sites to decrease YAP/TAZ activity. This function of c-JUN is independent of its heterodimeric AP-1 partner c-FOS demonstrating that it is independent of the canonical AP-1 function to induce target gene expression. Since c-JUN is itself by YAP/TAZ, our work identifies a negative feedback loop that buffers YAP/TAZ activity at joint sites. This negative feedback loop needs to get disrupted in liver cancer to unlock the full oncogenic potential of YAP/TAZ.

Our results thus demonstrate an additional layer of control for the important interplay of YAP/TAZ and AP-1.

## Introduction

The transcriptional coactivators YAP/TAZ are the critical downstream regulators of the Hippo pathway that regulate gene expression in response to changes in pathway activity, mainly by binding to TEAD transcription factors (Dong et al. 2007; Zhao et al. 2007). Uncontrolled transcriptional output of YAP/TAZ can lead to rapid induction of aggressive tumor growth, e.g. in the liver (Zhou et al. 2009; Yimlamai et al. 2014; Moya et al. 2019; Wu et al. 2022). For this reason, it is imperative for an organism to ensure tight and well-orchestrated control over YAP/TAZ to protect the body from the fatal consequences of their derailed activity (Driskill and Pan 2021).

AP-1 is a dimeric basic leucine zipper (bZIP) transcription factor complex with JUN and FOS proteins being the most abundant members of this family (Eferl et al. 2003). Unlike FOS, JUN can also form homodimers, but in the cell, JUN preferentially forms heterodimers with members of the FOS family, which act as potent transcriptional activators (Vogt 2002). Previous studies identified substantial co-occupancy of YAP/TAZ and AP-1 at genomic sites, and they demonstrated that JUN/FOS heterodimers cooperate with the YAP/TAZ and TEAD transcription factors to drive YAP/TAZ target gene expression (Shao et al. 2014; Zanconato et al. 2015; Koo et al. 2020).

A complete understanding of all YAP/TAZ control mechanisms may thus provide a basis for novel cancer therapies. On the other hand - due to the essential roles of these coactivators in regeneration (Leach et al. 2017; Elster and von Eyss 2020) - targeted modulation of YAP/TAZ could additionally be used to improve this process.

In this article, we now elucidate a negative feedback mechanism in which high YAP activity is restrained by the recruitment of homodimeric JUN::JUN/NCOR1 repressor complexes and show that this noncanonical JUN function is part of a tumor suppressor mechanism in the liver.

## Results

### c-JUN antagonizes YAP5SA-mediated growth arrest

We observed that strong lentiviral overexpression of the hyperactive YAP5SA mutant in MCF10A cells leads to potently reduced growth of these (Fig. 1A,B). Here, we made use of this overexpression phenomenon to screen for genes that could – when overexpressed – suppress this phenotype. We used the genome-wide *Synergistic Activation Mediator* (SAM) library (Konermann et al. 2015) to identify genes that suppress this phenotype when overexpressed. MCF10A-SAM cells infected with a genome-wide SAM library were subsequently superinfected with YAP5SA. Cells were kept in culture for two weeks to allow outgrowth of cells expressing potential suppressors of YAP5SA (Fig. 1B). Our analyses identified numerous genes that were significantly enriched in this screen (Fig. 1C, Supplementary Table 1) and MYC was among the top hits, which was previously described as a Hippo pathway-independent suppressor of YAP (von Eyss et al. 2015; Croci et al. 2017). *JUN* (c-JUN) particularly piqued our interest because several studies proposed a cooperative behavior between YAP and AP-1 in cancer cells (Stein et al. 2015; Zanconato et al. 2015; Koo et al. 2020). Our data, however, would argue that c-JUN suppresses YAP function. For this reason, we followed up more closely on this interesting but seemingly counterintuitive finding. In our SAM screen, *JUN* was the only member of the JUN family that showed an enrichment of sgRNAs since *JUND* and *JUNB* sgRNAs were not enriched (Fig. 1D). We validated the results from our SAM screen using MCF10A-SAM cells infected with individual sgRNAs (Fig. 1E). Like in the SAM screen, sgJUN#1 was more potent than sgJUN#2 in terms of its ability to rescue cell growth, which correlated with JUN expression on protein level (Fig. 1F).

**Fig. 1:**
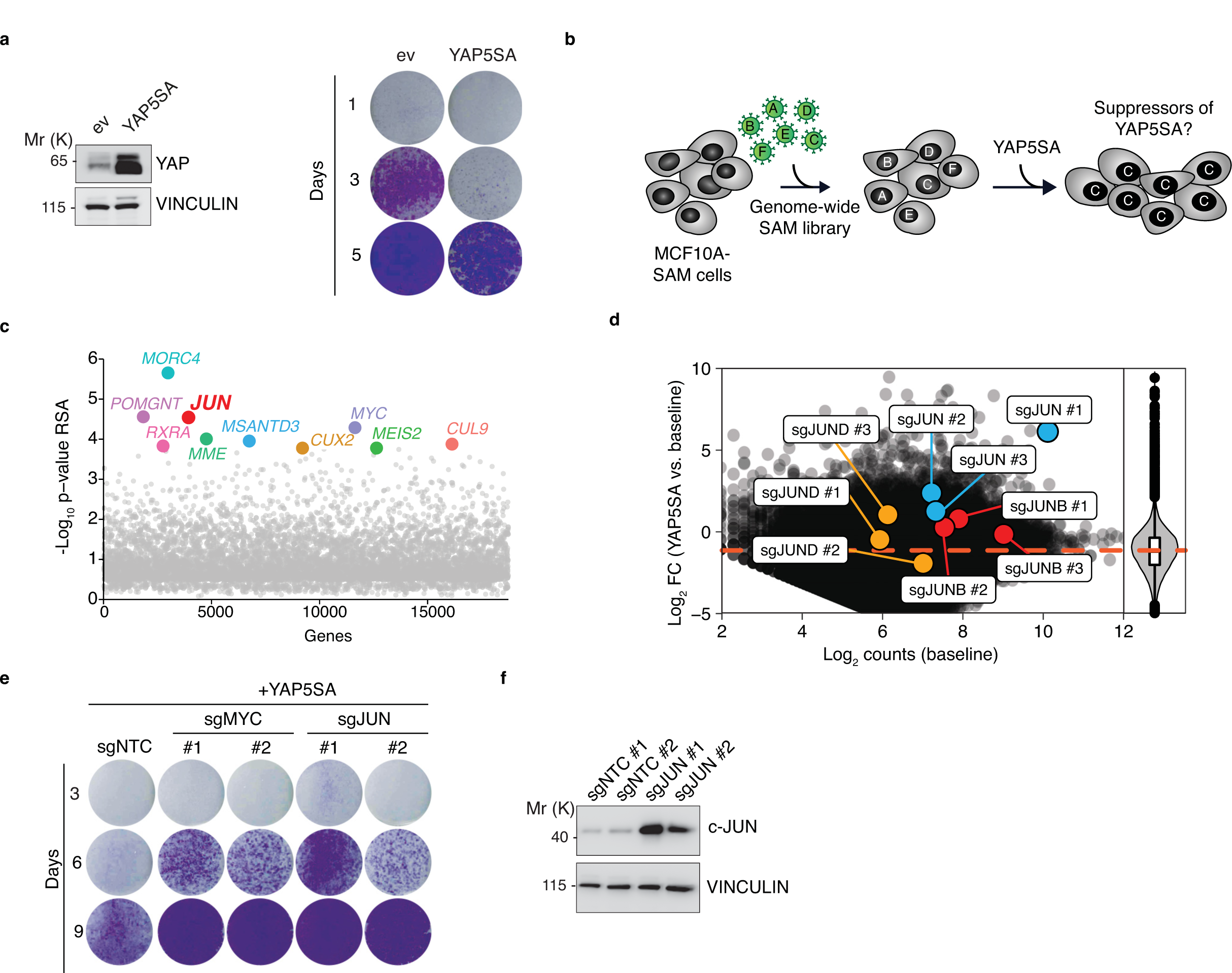
c-JUN antagonizes YAP5SA-mediated growth arrest. (a) Immunoblot (left) and crystal violet (right) from MCF10A cells infected with the indicated lentiviral vectors. Cells were stained with crystal violet after 1, 3, and 5 days post-infection. (b) Schematic illustrating the SAM screen to identify suppressors of YAP5SA-mediated growth arrest. (c) Summary of SAM screen for enrichment of sgRNAs (14 days after YAP5SA overexpression / baseline) targeting all human genes. The redundant siRNA algorithm (RSA) was used to integrate all sgRNAs targeting one gene and to infer statistical significance of enrichment. (d) MA plot illustrating the distribution of sgRNAs targeting JUN family members. The violin plot (right) illustrates the Log2 fold changes (YAP5SA vs. baseline) for all sgRNAs that are plotted in the MA plot. (e) Crystal violet of MCF10A SAM cells expressing individual sgRNAs which were superinfected with YAP5SA and stained after 3, 6, and 9 days of YAP5SA infection. sgNTC=Non-targeting control. (f) Immunoblot from MCF10A SAM cells infected with the indicated JUN and control sgRNAs. sgNTC=Non-targeting control.

### c-JUN interferes with induction of a large fraction of YAP target genes

We next analyzed the transcriptome of JUN-overexpressing (sgJUN#1) MCF10A-SAM cells. Gene set enrichment analysis (GSEA) showed that YAP target genes were the most potently downregulated gene set after JUN overexpression (Fig. 2A,B). These results could be corroborated with cDNA-mediated expression of JUN since a mild JUN overexpression led to induction of the AP-1 target gene *IL1B* and to potent downregulation of *ANKRD1* and *THBS1* mRNA expression (Fig. 1C,D), which are both part of the Cordenonsi YAP gene set. In addition, JUN was able to suppress YAP-dependent induction of the *ANKRD1* promoter in a reporter assay (Supplemental Fig. S1A,B). To investigate a potential involvement of the Hippo pathway in this, we used an MCF10A cell line (iYAP5SA) expressing the Hippo-insensitive YAP5SA allele (Zhao et al. 2007) in a doxycycline-dependent manner at sub-endogenous levels (von Eyss et al. 2015). First, we performed independent RNA-Sequencing (RNA-Seq) experiments applying a short doxycycline treatment to enrich for direct YAP target genes and limit secondary effects (Supplemental Fig. S2A,B, Supplementary Table 2). We then inferred the Top-YAP-induced (TYI) genes (log2FC < 1, padj < 1e-4, n=434 genes, Supplementary Table 2) from this experiment for our consecutive analyses. Next, iYAP5SA cells were infected with JUN expression vector or a vector control (Fig. 2E). Consistent with previous reports (Maglic et al. 2018), YAP was able to induce JUN expression at the protein level, similar to JUN overexpression by cDNA (Fig. 2E). Subsequent RNA-Seq and unsupervised clustering analyses (Fig. 2F,G) revealed three main clusters (Supplementary Table 3) within TYI genes: cluster 3 genes mainly consisted of known AP-1-induced target genes, cluster 1 and cluster 2 contained well-established direct YAP/TAZ target genes (Fig. 2G). Whereas JUN overexpression had only a very weak or no effect on cluster 2 genes, YAP-dependent induction of cluster 1 genes was completely blunted by JUN overexpression (Fig. 2F-I). Next, we performed SLAM-Seq experiments (Fig. 2J) in conjunction with acute JUN protein depletion, as described previously for MYC(Muhar et al. 2018). MCF10A JUN knockout cells were reconstituted with a JUN allele fused to an V5-tagged auxin-inducible degron (JUN-AID-V5) which was rapidly degraded after one hour of indole-3-acetic acid (IAA) addition (Fig. 2K, Supplemental Fig. S3A,B). Based on the T-to-C conversions in SLAM-Seq that occur in *de novo* synthesized mRNAs at the 4-thiouridine (4sU) sites, one can distinguish between changes at the steady-state level, and changes in *de novo* synthesized mRNA during the 4sU pulse/JUN degradation phase (Fig. 2L). Whereas there were no significant changes in the fraction of total mRNAs of YAP/TAZ target genes (Supplementary Table 4), these target genes were significantly induced in the fraction of *de novo* mRNAs (Fig. 2L).

**Fig. 2:**
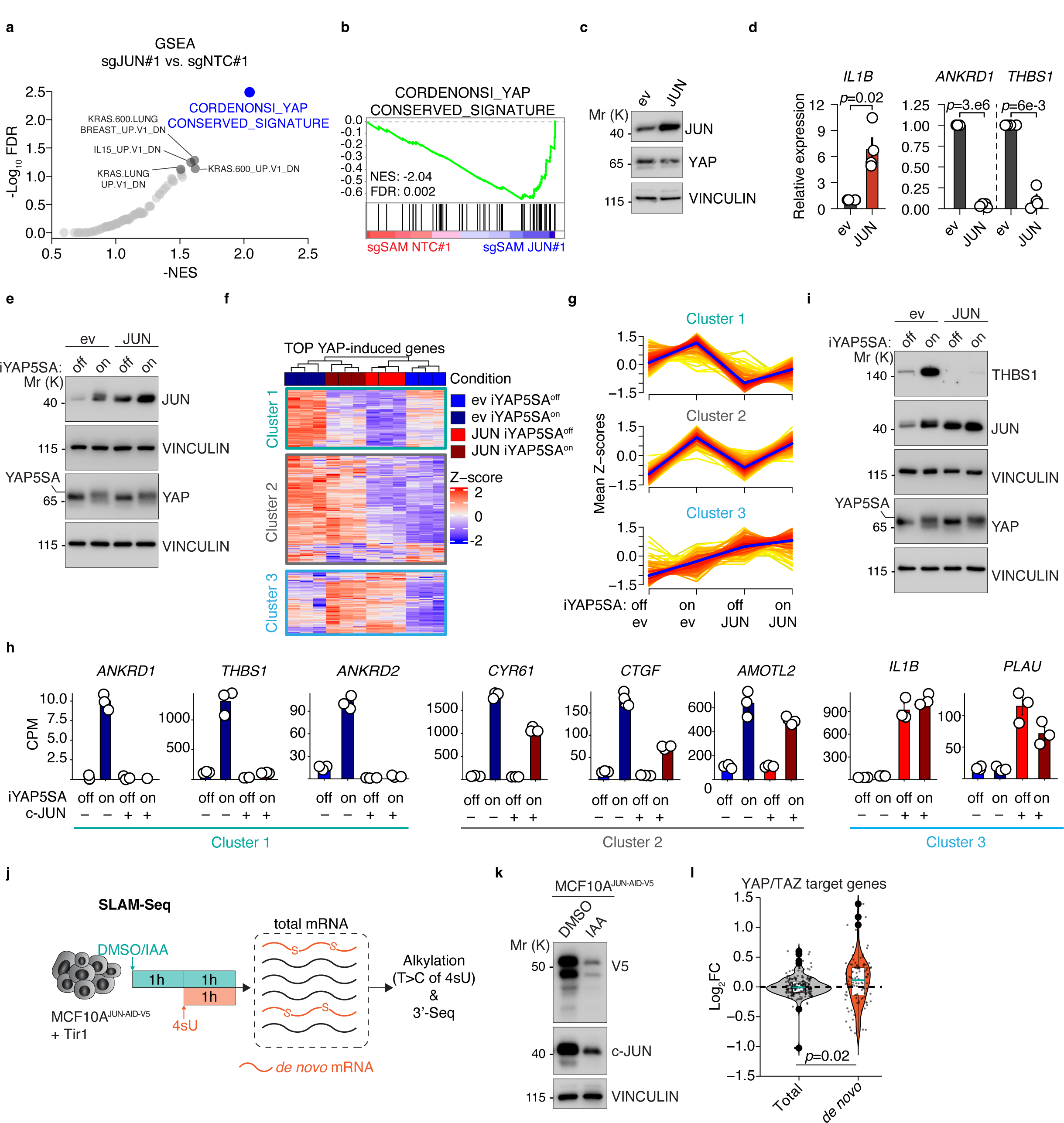
c-JUN interferes with YAP target gene expression. (a) GSEA summary of downregulated gene sets from RNA-Seq data comparing sgJUN#1 vs. sgNTC#1 cells. NES=normalized enrichment score, FDR=false discovery rate (b) GSEA enrichment plot for the Cordenonsi gene signature as the most strongly downregulated gene set in sgJUN#1. NES=normalized enrichment score, FDR=false discovery rate (c) Immunoblot of MCF10A SAM cells infected with JUN cDNA vectors or an empty vector control (ev). (d) qRT-PCR analysis of MCF10A SAM cells after JUN cDNA overexpression. Summary of four biological replicates. Welch T-test. (e) Immunoblot analysis of iYAP5SA cells infected with JUN overexpression vectors or an empty vector control (ev). YAP5SA was induced for 16 h and subsequently analyzed. (f) Heatmap of TOP YAP-induced genes (Z-score normalized, n=434 genes) in the indicated experimental conditions in iYAP5SA cells, analyzed by RNA-Seq and subsequent unsupervised clustering with three clusters. n=3 biological replicates per experimental group. (g) Summary of mean Z-scores (per experimental group) for all TOP YAP-induced genes to illustrate the expression changes in the four experimental groups. (h) Barplots illustrating the expression of representative genes for each cluster in the RNA-Seq experiment. CPM=mapped counts per million. (i) Immunoblot analysis of iYAP5SA cells that were infected with JUN overexpression vectors or an empty vector control (ev). YAP5SA was induced for 16 h and subsequently analyzed. (j) Schematic of the SLAM-Seq approach that combines acute JUN depletion via the auxin system with metabolic labeling of *de novo* mRNA transcripts using 4-thiouridine (4sU) in MCF10A^JUN-AID-V5^ cells. IAA=indole-3-acetic acid. (k) Immunoblot from MCF10A^JUN-AID-V5^ cells 2 hours after indole-3-acetic acid (IAA) addition to induce JUN degradation. (l) Violin plots of YAP/TAZ target genes that illustrate the gene expression changes upon acute JUN depletion for the total mRNA pool (left) versus the de novo transcribed pool (right). Two-sided Wilcox test. Source data are provided as a Source Data file.

### c-JUN is recruited by YAP and interferes with YAP/TEAD recruitment

Based on the results of the SLAM-Seq, we hypothesized that JUN directly interferes with YAP/TEAD on chromatin. To identify the sites bound by YAP/TEAD, as wells as JUN/FOS, we performed CUT&RUN experiments from MCF10A constitutively overexpressing YAP5SA (Fig. 3A-C) as well as from iYAP5SA infected with JUN expression vectors (Fig. 3D-I). Consistent with previous results in cancer cell lines, JUN and c-FOS bound to many YAP/TEAD target sites (joint Y/T AP-1), but they also bound to exclusive sites (AP-1 only), where no YAP/TEAD could be detected (Fig. 3A,B). AP-1-related motifs were strongly enriched in both YAP and TEAD1 peaks (Fig. 3C). We next investigated how JUN binding is affected by acute YAP induction, and *vice versa*, how JUN overexpression impacts on YAP binding. It should be mentioned here that this analysis is made more difficult by the fact that JUN itself is induced by YAP, so that one can also observe effects that simply occur due to increased JUN protein levels and are not necessarily mediated by recruitment of YAP to genomic sites.

**Fig. 3:**
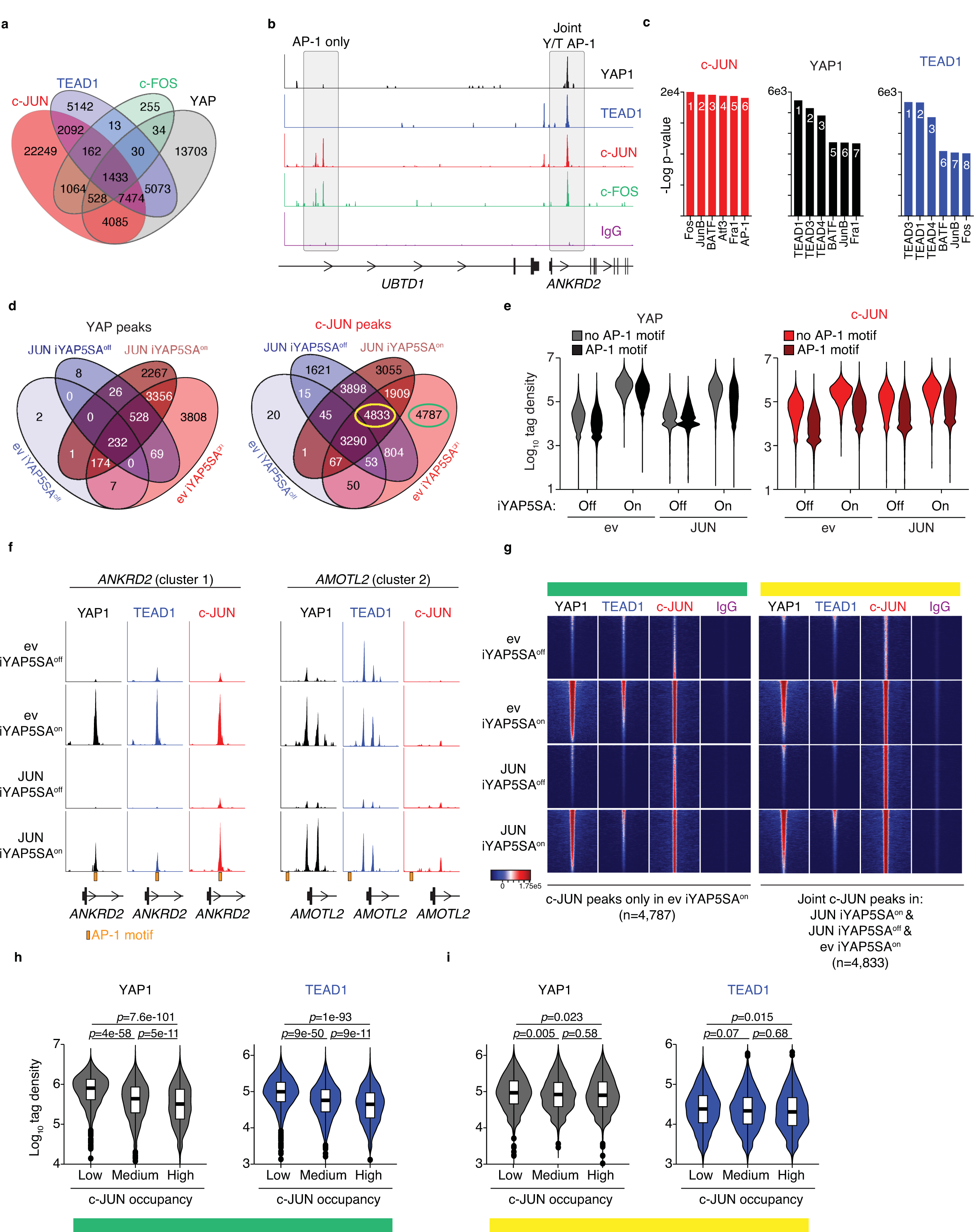
c-JUN is recruited by YAP to genomic sites. (a) Venn diagram for CUT&RUN peaks in YAP5SA-overexpressing MCF10A cells. (b) Representative CUT&RUN sequencing tracks that illustrate binding of AP-1 only (JUN and FOS, left) and joint YAP/TEAD AP-1 (Y/T AP-1, right) sites. (c) Motif enrichment analysis for all CUT&RUN peaks from JUN, YAP, TEAD1. The numbers indicate the rank in the motif analysis. (d) Venn diagram for YAP and JUN CUT&RUN peaks in MCF10A-iYAP5SA cells infected with a JUN expression construct or empty vector control (ev). One day prior to CUT&RUN, cells were treated with either EtOH as a solvent control (YAP5SA^off^) or doxycycline (YAP5SA^on^). (e) Count distribution within 200 bp windows of 16,307 joint YAP/TEAD peaks from (a) stratified based on the presence of an AP-1 motif in the peak. (f) Representative CUT&RUN sequencing tracks that illustrate the binding to *ANKRD2* and *AMOTL2*. AP-1 DNA binding motifs are highlighted with an orange box. (g) CUT&RUN heatmaps for binding of YAP1, TEAD1 and JUN to 4,787 JUN peaks that occur in a YAP-dependent manner (exclusively in the ev YAP5SA^on^ condition, highlighted in green). A control peak set of similar size with 4,833 peaks is highlighted in yellow. See also (d) for the peaks that were used here. All heatmaps were sorted based on the YAP signal in the ev YAP5SA^on^ situation. (h,i) Violin plots for tag densities of YAP and TEAD CUT&RUN data in 4,787 YAP-dependent JUN peaks (h, green) and a control peak set with 4,833 peaks (i, yellow). The tag densities were stratified based on the JUN signal (low, medium, high) in the JUN iYAP5SA^off^ condition. Two-sided Wilcox test with Benjamini-Hochberg correction.

Whereas JUN overexpression in uninduced conditions (JUN iYAP5SA^off^) led to 1,621 additional JUN sites (Fig. 3D), YAP induction in empty vector-infected cells (ev iYAP5SA^on^) led to 4,787 unique sites, arguing that these YAP-dependent sites cannot be simply explained by increased JUN protein levels. YAP induction itself led to a potent recruitment of YAP and JUN at joint YAP/TEAD sites (n=13,975 sites, Fig. 3A,E). While recruitment of YAP was largely unaffected by the presence of an AP-1 binding motif at these sites, recruitment of JUN was more pronounced in peaks with an AP-1 motif (Fig. 3E,F). This indicates that DNA-binding of JUN contributes to YAP-dependent recruitment. We next analysed in more detail the JUN peaks that occurred in a strictly YAP-dependent manner (n=4,787 peaks), together with a control peak set of similar size (n=4,833 peaks) that also included peaks in which JUN overexpression alone was sufficient to induce recruitment, suggesting that these sites are rather influenced by JUN expression rather than recruitment by YAP. For the YAP-dependent JUN peaks, it became apparent that JUN negatively impacts YAP/TEAD1 recruitment since high JUN binding (in empty vector or JUN overexpression conditions) was associated with strongly decreased YAP/TEAD1 recruitment after YAP induction, which was not observed for the control peak set (Fig. 3G-I).

### Canonical AP-1 function and YAP inhibition are distinct JUN properties

T-5224 is an inhibitor that was designed to interfere with AP-1 function by binding AP-1’s basic region required for DNA binding (Tsuchida et al. 2004; Aikawa et al. 2008). Furthermore, T-5224 can interfere with growth of YAP-dependent liver cancers (Koo et al. 2020). We tested whether T-5224 would specifically impact JUN-dependent YAP targets. Despite JUN induction on protein level (Fig. 4A), T-5224 reduced the expression of canonical AP-1 targets such as *IL1B* or *PLAU* (Fig. 4a,b). In addition, cluster 1 genes were potently downregulated, whereas the expression of cluster 2 genes barely changed (Fig. 4B). One reason for T-5224’s ability to downregulate cluster 1 genes could be the fact that FOS can enhance expression of YAP target genes (Koo et al. 2020), but that alternative JUN-containing AP-1 complexes exist, which in turn mediate the repression of YAP target genes. Thus, the ability of JUN to limit the transcriptional activity of YAP appears to be distinct from its canonical function in AP-1, i.e. induction of JUN/FOS target genes such as *IL1B*.

**Fig. 4:**
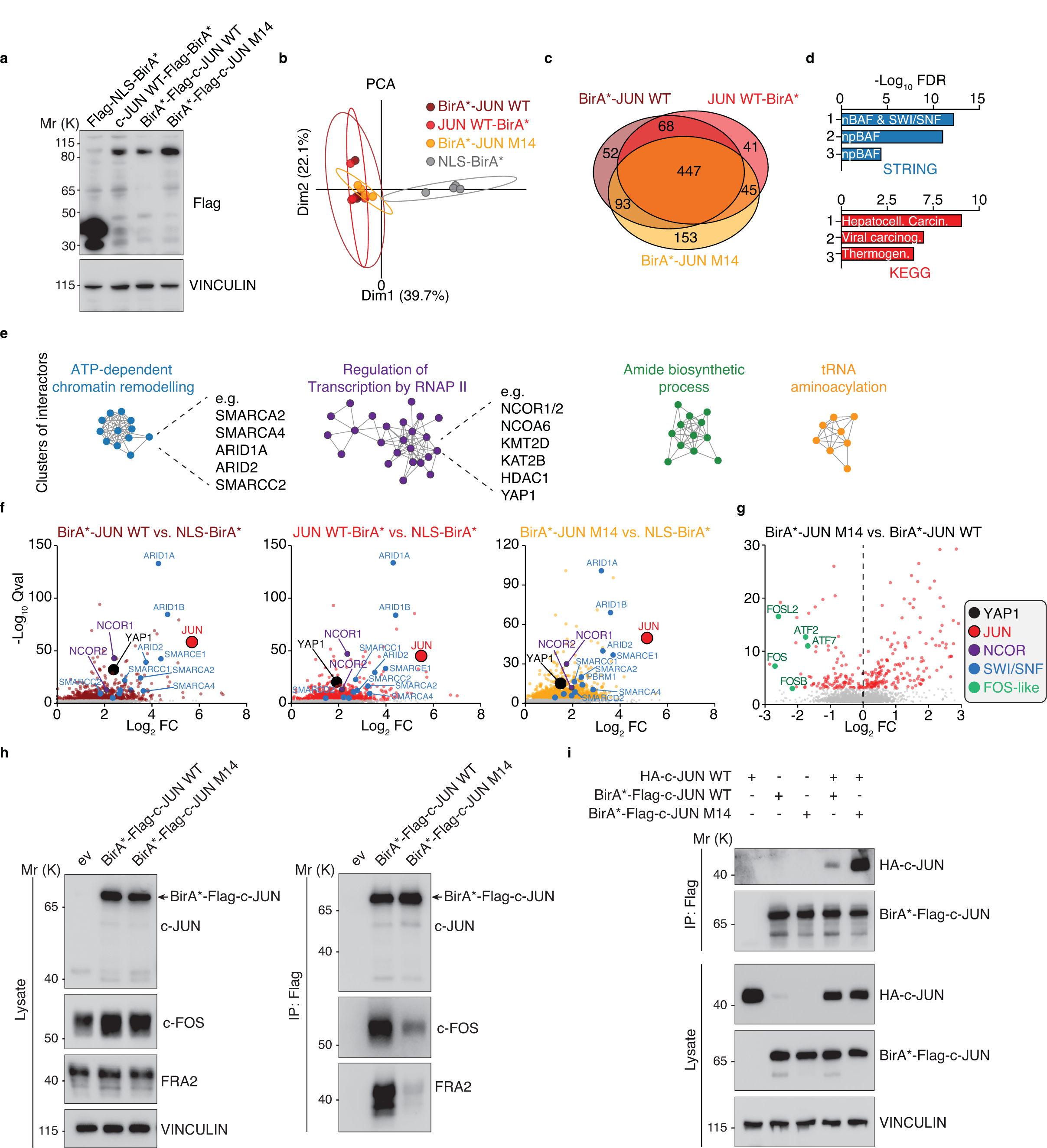
Canonical AP-1 function and YAP inhibition are distinct JUN properties. (a) Immunoblot of MCF10A cells that were treated with 100 µM of the AP-1 inhibitor T-5224 for 20h. (b) qRT-PCR of MCF10A cells that were treated with 100 µM of the AP-1 inhibitor T-5224 for 20h. n=3 biological replicates per group. Two-sided Welch test with Benjamini-Hochberg correction. (c) Schematic illustrating the mutations of JUN M14 and JUN I10. A blue box in the JUN::FOS crystal structure (left) marks the localization of the mutation in the JUN leucine zipper. The point mutations are shown in red, and the localization of the four amino acid insertion in JUN I10 is indicated. (d) Immunoblots of MCF10A cells infected with the indicated JUN alleles or an empty vector control (ev). (e) qRT-PCR analysis of MCF10A cells infected with JUN WT, JUN M14 or empty vector control (ev). One-way ANOVA with Tukey HSD post-hoc test. (f) Venn diagram of CUT&RUN peaks of JUN WT and JUN M14. CUT&RUN was performed in MCF10A JUN KO cells that were either reconstituted with JUN WT or JUN M14. (g) CUT&RUN heatmaps for JUN WT and JUN M14 binding to 24,448 JUN peaks defined previously (see JUN Venn diagram Fig. 3d). (h) Violin plots for the tag densities of JUN WT and JUN M14 CUT&RUN data in 5,335 joint YAP/JUN or 10,468 JUN only peaks. Two-sided Wilcox test with Benjamini-Hochberg correction. (i) Representative CUT&RUN sequencing tracks that illustrate the binding of JUN WT and JUN M14 to *ANKRD2* and *IL1B*.

To test this, we set out to find JUN mutants that would allow us to separate these two different JUN functions. Previous work identified leucine zipper mutants of JUN that showed differential activities towards activating AP-1-dependent reporter genes, potentially due to different abilities to form specific AP-1 homo- and heterodimers (Smeal et al. 1989). Here, we tested two mutants: JUN M14 and JUN I10. The M14 mutant harbours three point mutations in the leucine zipper region and retains the ability to bind DNA, whereas the insertion of four amino acids into the leucine zipper region of the JUN I10 mutant abolishes DNA binding (Fig. 4C). JUN WT overexpression in MCF10A cells induced IL1B expression and downregulated THBS1 on protein level, while JUN I10 did neither affect induction of IL1B nor led to downregulation of THBS1 (Fig. 4D). JUN M14, however, was able to potently decrease THBS1 expression whereas IL1B and CYR61 remained largely unchanged (Fig. 4D,E). Since JUN M14 showed a clear difference in terms of canonical AP-1 functions versus repression of YAP target genes, we analysed this mutant in more detail, as it would allow us to separate these two JUN functions. CUT&RUN experiments for JUN in JUN KO cells that were either reconstituted with JUN WT, JUN M14 or a vector control showed that JUN M14 had a similar binding behaviour as JUN WT (Fig. 4F,G). Furthermore, we were not able to identify any consistent differences in their binding behaviour towards joint YAP/JUN targets or JUN only targets, even though JUN M14 tended to show a stronger signal (Fig. 4H,I) which could be simply due to higher expression levels (Fig. 4D). Thus, JUN M14 is unable to activate canonical AP-1 target genes but retains the ability to bind to genomic targets and interfere with YAP function.

### JUN M14 shares a similar interactome with JUN WT but is deficient in heterodimerization with FOS

To identify critical mediators of JUN-dependent repression, we performed BioID experiments in MCF10A JUN KO cells that were reconstituted with different Flag-BirA* JUN fusion proteins or nuclear localization signal (NLS)-Flag-BirA* control (Fig. 5A). In the mass spectrometry analysis (see Supplementary Table 5 for a list of all interactors) all JUN proteins showed a strikingly similar interactome in the principal component analysis or the overlap of all interactors (Fig. 5B,C). Here, a strong enrichment for components of SWI/SNF complexes and proteins involved in hepatocellular carcinogenesis could be observed (Fig. 5D). Cluster analysis of interactors showed that ATP-dependent chromatin remodelling components (e.g. ARID1A and SMARCA2/4) and regulators of RNA polymerase (RNAP) II (e.g. NCOR1/2) were high confidence interactors of JUN WT and JUN M14 (Fig. 5E,F). We also identified YAP as a JUN WT/M14 interactor, confirming our hypothesis that JUN is recruited to shared YAP/JUN sites via a protein interaction with YAP. When comparing JUN WT with JUN M14, all FOS-like proteins (e.g. FOS, FOSB, FOSL2) showed at least a 4-fold reduced labelling efficiency in JUN M14 (Fig. 5G). This was confirmed by co-immunoprecipitation, as FOS and FRA2 were significantly reduced in Flag precipitates from BirA*-Flag JUN M14 compared with BirA*-Flag JUN WT (Fig. 5H). On the other hand, JUN M14 was still able to homodimerize with JUN WT, which even tended to be more pronounced in JUN M14 compared with JUN WT (Fig. 5I). This suggests that the impaired function of JUN M14 to induce canonical AP-1 targets is due to its reduced interaction with FOS proteins, whereas the effect on YAP targets could potentially be mediated by JUN::JUN homodimers.

**Fig. 5:**
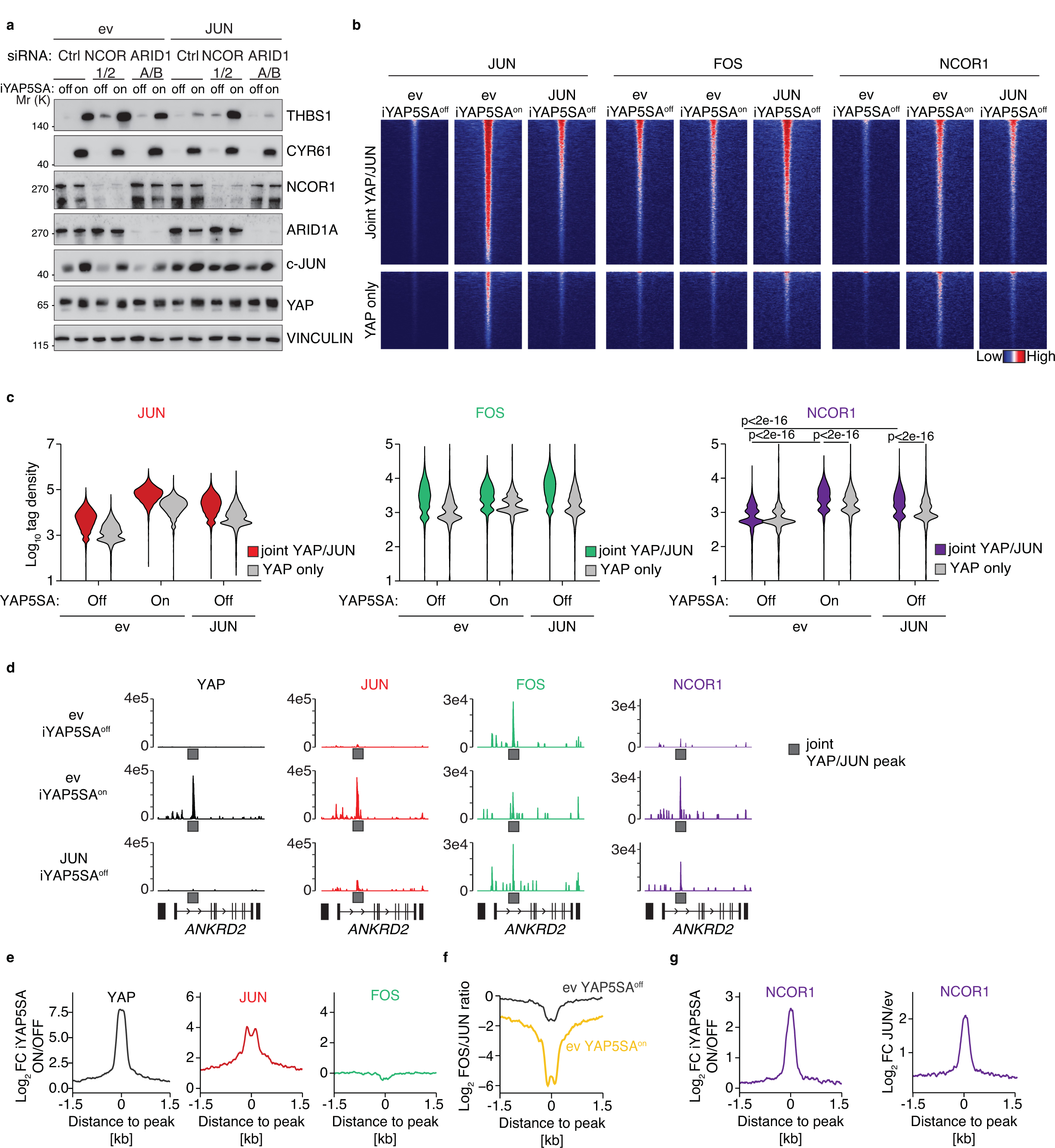
JUN M14 shares a similar interactome with JUN WT. (a) Immunoblot of MCF10A JUN KO cells reconstituted with the indicated BirA*-Flag alleles. (b) Principal component analysis (PCA) for the BioID experiment. (c) Venn diagram for the high confidence interactors (q-val<0.001, Log2FC >1) of the indicated BirA*-Flag JUN alleles identified by BioID. (d) Functional annotation of the 447 common JUN interactors using the STRING or KEGG database. FDR=false discovery rate (e) Clustering analysis of 447 common JUN interactors. (f) Volcano plots for the enrichment of the different BirA* JUN fusion proteins versus the NLS-BirA* control. (g) Volcano plots for the enrichment of the BirA*-Flag JUN WT versus BirA*-Flag JUN M14. (h) Co-immunoprecipitation experiments from MCF10A JUN KO cells reconstituted with the indicated alleles. BirA* fusion proteins were immunoprecipitated with Flag, and precipitates were assayed for endogenous FOS and FRA2. (i) Exogeneous co-immunoprecipitation experiments from 293T cells transfected with the indicated expression constructs. BirA* fusion proteins were immunoprecipitated with Flag, and precipitates were assayed for HA-JUN WT.

### NCOR1/2 are required for JUN’s ability to repress YAP target genes

To identify repressor proteins that could mediate the repressor function of putative JUN::JUN homodimers, we focused on the SWI/SNF component ARID1A/B and the corepressor protein NCOR1/2 since both can behave as repressors of transcription and were potently enriched for all JUN proteins (Fig. 5B). To this end, we depleted ARID1A/B and NCOR1/2 in iYAP5SA cells overexpressing JUN WT by siRNA. Whereas ARID1A/B-depleted cells behaved like control-depleted cells, NCOR1/2 depletion was able to completely restore THBS1 in JUN-overexpressing cells under YAP5SA-induced conditions (Fig. 6A, Supplemental Fig. S4). Since these experiments identify NCOR1/2 as a critical JUN-interacting protein for its ability to repress YAP target genes, we next tested whether YAP and/or JUN are able to recruit NCOR1/2 to common genomic sites, and whether the binding behavior is consistent with recruitment by JUN::JUN homodimers.

**Fig. 6:**
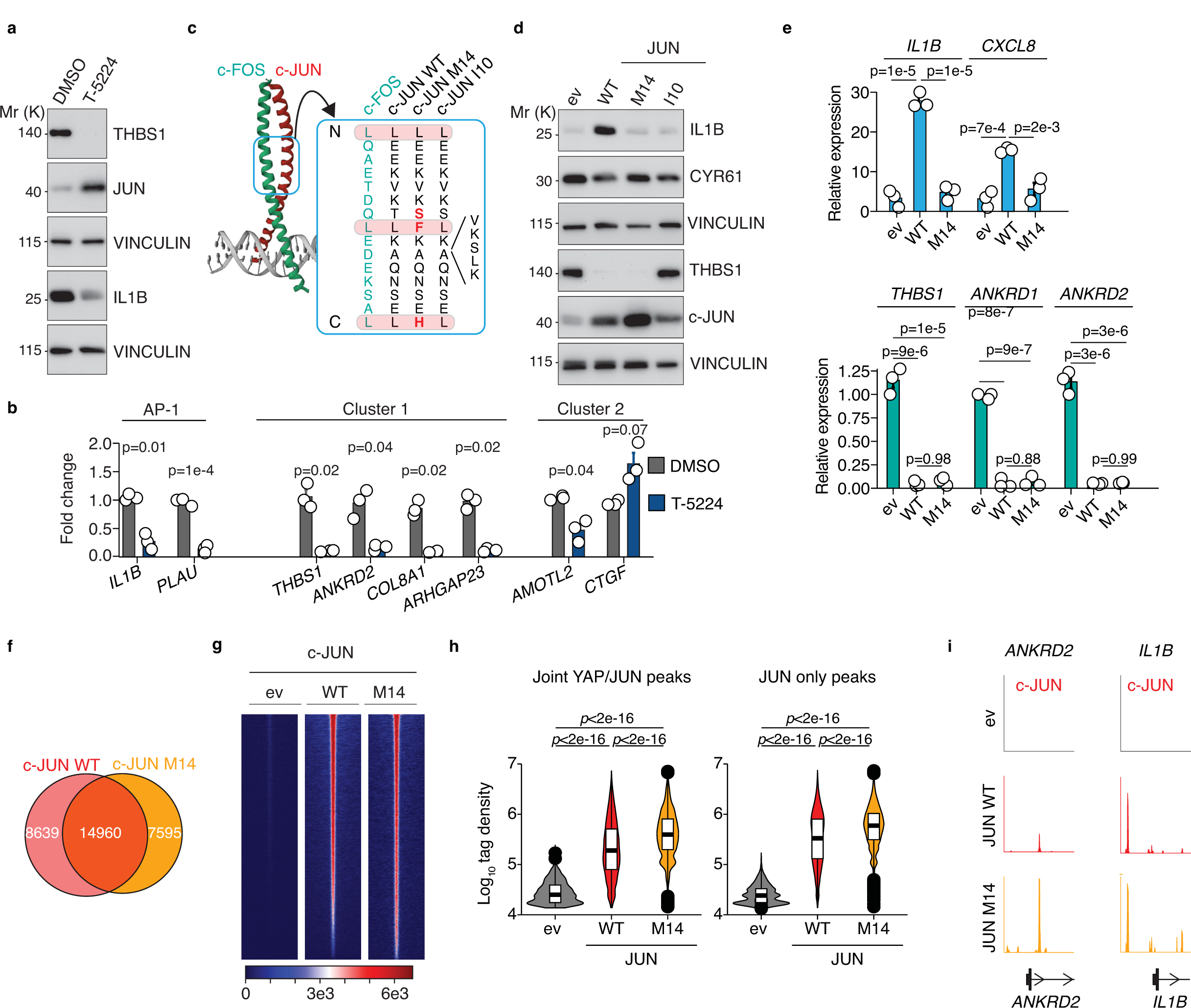
JUN depends on NCOR1/2 to repress YAP target genes. (a) Immunoblot from iYAP5SA MCF10A cells infected with JUN WT or empty vector control (ev) which were transfected with the indicated siRNAs. YAP5SA expression was induced for circa 20 hours with doxycycline or cells were treated with EtOH as solvent control. (b) CUT&RUN heatmaps for JUN, YAP, and NCOR1 to 5,335 joint YAP/JUN peaks and 10,468 JUN only peaks, respectively. Three different conditons in iYAP5SA MCF10A cells were analysed: empty vector-infected uninduced (eviYAP5SA^off^), empty vector-infected YAP5SA-induced (ev iYAP5SA^on^), and JUN overexpression uninduced (JUN iYAP5SA^off^). (c) Violin plots for the tag densities of JUN, YAP, and NCOR1 at the indicated peaks in a 200 bp window centred on the peak summit for then same peaks and conditions as in a. Two-sided Wilcox test. (d) Representative CUT&RUN sequencing tracks for the indicated proteins and conditions at the *ANKRD2* gene. (e) YAP-dependent recruitment (ev iYAP5SA^on^ vs. ev iYAP5SA^off^) of the indicated proteins to 5,335 joint YAP/JUN peaks. (f) FOS/JUN signal ratios at the 5,335 joint YAP/JUN peaks for the ev iYAP5SA^off^ and ev iYAP5SA^on^ conditions. (g) JUN-dependent recruitment (JUN iYAP5SA^off^ vs. ev iYAP5SA^off^) of NCOR1 to 5,335 joint YAP/JUN peaks.

We performed CUT&RUN experiments for JUN, FOS, and NCOR1 in iYAP5SA under three conditions: control condition, YAP5SA-induced, and JUN overexpression. As before, YAP5SA induction led to a strong recruitment (∼10-fold) of JUN to joint YAP/JUN sites, whereas JUN overexpression achieved a milder recruitment of JUN to the same sites (Fig. 6B-D). Despite potent JUN recruitment, FOS remained largely unchanged upon YAP5SA induction or JUN overexpression (Fig. 6B-D). In contrast, NCOR1 binding was significantly increased by both YAP and JUN, and NCOR1 binding was stronger at joint YAP/JUN sites compared to YAP only sites (Fig. 6B-D). The differential analysis of YAP5SA-induced vs. uninduced corroborated that YAP5SA can potently induce binding of YAP and JUN to joint YAP/JUN sites, whereas FOS binding rather tended to be slightly reduced (Fig. 6E). Thus, YAP induction leads to a sharp drop in the FOS/JUN ratio at joint YAP/JUN sites which is consistent with recruitment of JUN::JUN homodimers that would not lead to elevated FOS binding to these sites (Fig. 6F).

On the other hand, YAP5SA induction or JUN overexpression itself was sufficient to induce NCOR1 recruitment, suggesting that YAP-dependent recruitment of NCOR1 may be mediated by JUN (Fig. 6G).

### JUN suppresses YAP-dependent liver cancers

As described before (Fig. 2E), and shown by others (Maglic et al. 2018), YAP can induce JUN expression on protein level. This suggests that JUN is part of a negative feedback loop to restrain expression of cluster 1 genes (Fig. 7A). We thus wondered whether this negative feedback loop gets disrupted in cancer to unleash YAP’s full transcriptional potential. We therefore performed a differential analysis using genes from cluster 1 and cluster 2 as readout of YAP transcriptional activity to identify scenarios in which specifically this feedback loop is disrupted (Fig. 7A,B). That way, one should be able to discriminate between a general increase in YAP activity (e.g. by YAP overexpression), and a potential disruption of the feedback loop since the latter would specifically affect cluster 1 genes. To this end, 19 different cancer entities with 7,458 patients in TCGA data sets were analyzed for a specific survival benefit when comparing expression cluster of 1 vs. cluster 2 genes (Fig. 7B). In this analysis, hepatocellular carcinoma (HCC) stood out, as in particular the survival probability of patients with low expression of cluster 1 genes was strongly increased, whereas the hazard ratio for cluster 2 genes was largely unaffected (Fig. 7B,C). To test whether JUN could suppress YAP-dependent liver cancer, we performed hydrodynamic tail vein injection (HDTVI) in conjunction with a sleeping beauty-based approach to stably express genes in the livers of C57BL/6J wildtype mice. YAP overexpression in combination with constitutively active myristoylated Akt (myr-AKT) leads to induction of hepatocellular carcinomas only a few weeks after HDTVI (Yamamoto et al. 2017). To investigate how JUN affects the oncogenic potential of YAP in this context, we co-overexpressed myr-AKT with the following constructs (Fig. 7D): GFP-IRES-YAP5SA (Ctrl), JUN WT-IRES-YAP5SA (JUN WT), and JUN M14-IRES-YAP5SA (JUN M14). Six weeks after HDTVI, the mice were sacrificed, and the livers were analyzed (Fig. 7E-I). The livers of the YAP5SA condition were strongly enlarged and showed numerous macroscopically visible tumor nodules whereas the JUN WT as well as the JUN M14 livers appeared largely normal (Fig. 7E). The liver to body weight ratios of YAP5SA-JUN WT and YAP5SA-JUN M14 animals were comparable to wildtype mice, while it was significantly elevated in YAP5SA animals (Fig. 7F). YAP5SA liver cancers showed a multifocal tumor growth with large tumor lesions whereas those lesions were barely detectable in YAP5SA-JUN WT/M14 livers, demonstrating that JUN’s repressive effect on YAP activity potently interferes with liver tumor growth (Fig 7G,H). On sections, all tumors were positive for HA-myr-AKT, nuclear YAP, and YAP5SA-JUN WT/M14 tumors showed strong JUN expression, demonstrating that the proteins were stably expressed as expected (Fig. 7I).

**Fig. 7:**
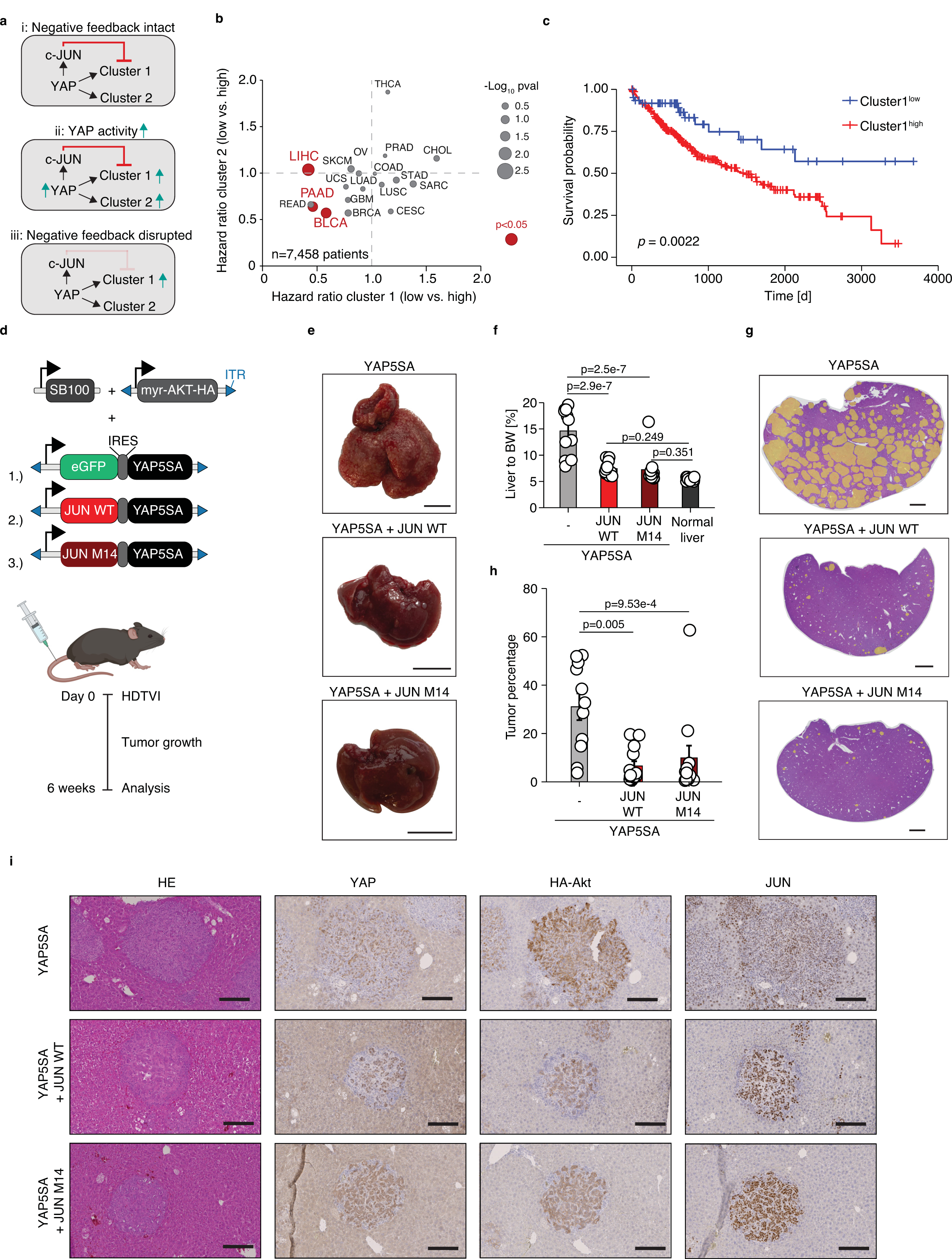
JUN represses YAP-dependent liver cancer growth. (a) Rationale to identify conditions in which specifically the negative feedback between JUN and YAP is lost. (b) Differential survival analysis in TCGA data sets for 7,458 cancer patients in 19 different cancer types. Per cancer type, each patient was analyzed for expression levels of cluster 1 and cluster 2 genes, respectively. Patients were stratified for low and high expression of both clusters, and the survival probability (hazard ratio) per cancer type was plotted against each other in both strata. Cancers with significant survival differences (p<0.05) for cluster 1 genes are highlighted in red. (c) Kaplan Meier curve for HCC patients that were stratified for expression of cluster 1 genes. Log-rank test. (d) Schematic for the liver cancer model by hydrodynamic tail vein injection (HDTVI). SB=Sleeping beauty transposase, ITR=inverted terminal repeat. (e) Representative Photos of mouse livers six weeks after HDTVI. Scale bar=1 cm. (f) Liver-to-body weight ratios for all mice six weeks after HDTVI. One-way ANOVA with Tukey HSD post hoc test. (g) H&E section of HDTVI liver tumors. Tumor nodules are colored in yellow. Scale bar: 2 cm. (h) Quantification of tumor areas (tumor area/ liver area) performed as in h. One-way ANOVA with Tukey HSD post hoc test. YAP5SA only (n=11), YAP5SA + JUN WT (n=13), YAP5SA + JUN M14 (n=12). (i) Representative immunohistochemical analyses of HDTVI liver tumors.

## Discussion

Numerous ChIP-Seq studies in cancer cells have documented a substantial overlap of YAP/TAZ, TEAD and AP-1 binding in the genome (Stein et al. 2015; Zanconato et al. 2015), and JUN/FOS AP-1 complexes are required for full transcriptional activity of YAP/TAZ (Shao et al. 2014). Our work now identifies an unexpected role of JUN::JUN homodimers whose function is to buffer unrestrained YAP/TAZ for a large fraction of YAP/TAZ target genes.

Most likely, JUN::JUN homodimers, together with its associated corepressor complexes, get recruited by protein interactions between YAP and JUN, and these interactions are further stabilized by DNA-binding of homodimers to AP-1 DNA sites. These sites seem to be pre-occupied by activating JUN::FOS heterodimers, so that JUN::JUN recruitment defines an AP-1 switch: from activating JUN::FOS to repressive JUN::JUN. Currently, it is unclear how the regulation of this switch is orchestrated: it cannot be simply explained by different ratios of JUN and FOS since JUN overexpression does not lead to a general switch of AP-1 to repressor because it still induces canonical AP-1 targets, such as *IL1B*. It is thus conceivable that YAP locally recruits additional proteins that mediate this switch, but this needs to be investigated in the future. Hepatocellular carcinoma is certainly one of the best characterized tumor entities with respect to the effects of uncontrolled YAP/TAZ activity. We now identify JUN as a component of a negative feedback mechanism with tumor suppressive properties to protect the organism from oncogenic YAP levels. At first glance, this finding seems counterintuitive since deletion of JUN leads to increased survival in a chemically induced liver cancer model (Eferl et al. 2003) which would suggest pro-oncogenic properties of JUN in liver cancer. However, deletion of JUN leads to removal both JUN activities: activating as well as repressive functions. It is therefore most likely context-dependent which of the two functions is more important for carcinogenesis, and this is certainly influenced by the driver mutations in each tumor. However, this dichotomy also implies that some of the described JUN knockout phenotypes may need to be reevaluated in this context, e.g. using JUN mutants such as the JUN-M14 mutant that can uncouple these two JUN functions.

Our analyses further suggest that therapies aiming to restore JUN’s repressive function in liver cancer could be beneficial in liver cancer. T-5224 would be a good candidate in this regard, since it interferes with activating AP-1 functions (Fig. 4B), and it shows a high efficacy in YAP-driven liver tumors (Koo et al. 2020).

In conclusion, our work defines a new layer of YAP/TAZ regulation in the context of the intricate AP-1 network which holds a great potential for developing novel cancer therapies.

## Methods

### Reagents

A list of reagents (antibodies, plasmids, oligonucleotides, siRNAs, chemicals, commercial kits) is provided in Supplementary Table 6.

### Mice

Animal experiments were conducted in accordance with the guidelines and regulations of the state government of Thuringia under the animal experiment license FLI-21-015. Male C57BL/6JRj mice for HDTVI experiments were obtained from Janvier Labs at 5 weeks of age. The mice were kept in individually ventilated cages (IVCs) under Specific Pathogen Free (SPF) conditions with a 12 h dark/12 h light cycles at 20°C and 55% relative humidity according to the directives of the 2010/63/EU and GV SOLAS.

### Mammalian cell culture

293T/LentiX cells (Takara Bio, # 632180) were cultured in DMEM (+GlutaMAX, Thermo Fisher Scientific) supplemented with 10% FBS (Thermo Fisher Scientific) and 1% penicillin-streptomycin (Sigma-Aldrich). MCF10A cells (kind gift from Martin Eilers, University of Würzburg, Germany) were cultured in DMEM/F12 (Thermo Fisher Scientific) supplemented with 5 % horse serum (Thermo Fisher Scientific), 1 % penicillin-streptomycin (Sigma-Aldrich), 10 µg/ml human insulin (Sigma-Aldrich), 0.5 µg/ml hydrocortisone (Sigma-Aldrich), 0.1 µg/ml cholera toxin (Sigma-Aldrich) and 20 ng/ml human EGF (Biomol). All cells were kept at 5% CO2, 95% relative humidity and 37°C and were regularly tested for mycoplasma contamination by PCR.

Transient transfections were carried out using polyethyleneimine (PEI-Max, Sigma-Aldrich) with Opti-MEM reduced serum medium (Thermo Fisher Scientific).

For siRNA transfections, the Lipofectamine RNAiMAX reagent (Thermo Fisher Scientific) was used. siRNAs were purchased from Dharmacon and are listed in Supplementary Table 6.

Expression of inducible YAP5SA mutant in MCF10A cells was performed by incubating the cells with 100 ng/ml doxycycline for 16 h prior to analysis. As control, the cells were treated with the same volume of ethanol.

For T-5224 treatment, the cells were incubated with 30 μM T-5224 (MedChemExpress) for 16 h before processing. Controls were treated with the same volume of DMSO.

### CRISPR-Cas9 KO

To generate MCF10A JUN knockout cells, a small guide RNA targeting the JUN exon (TCGTTCCTCCCGTGAGAG) was cloned into pX461 (a gift from Feng Zhang, Addgene #48140) in which the Cas9 nickase allele was substituted for the wild-type Cas9 allele. MCF10A cells were transfected using Lipofectamine 3000 (Thermo Fisher Scientific). After 48 h, GFP+ cells were isolated by flow cytometry and single cells were seeded in 96-well plates. JUN knockout clones were identified by Western blot. For knockout verification, genomic DNA was isolated and the regions flanking the sgRNA target site were amplified by PCR. The amplicons were cloned into pJET (Thermo Fisher Scientific) and analysed by Sanger sequencing, which revealed frame shift mutations leading to generation of premature stop codons.

### Acute JUN depletion via an auxin-inducible degron

JUN KO MCF10A cells were reconstituted with an auxin-tagged JUN allele (JUN-AID-V5). The cells were infected with a TIR1 overexpression construct, and JUN degradation was induced by adding 100 µM Indole-3-acetic acid (IAA, Sigma-Aldrich).

### Lentiviral transduction

For lentivirus production, LentiX cells were co-transfected with 10 μg psPAX2, 2.5 μg pMD2.G and 10 μg lentiviral vector using PEI-Max (Sigma-Aldrich).

Virus-containing supernatants were collected 24 h, 48 h, and 72 h post transfection and pooled. MCF10A cells were infected for 24 h using filtered viral supernatant diluted with culture medium and supplemented with 8 µg/µl protamine sulphate (Sigma-Aldrich). Selection of infected cells with antibiotics was performed 48 h after the infection.

### SAM screening

The genome-wide human CRISPR/Cas9 Synergistic Activation Mediator (SAM) sgRNA library (gift from Feng Zhang, Addgene #1000000057) was amplified as recommended, and balanced sgRNA library distribution was verified by NGS. Clonal MCF10A cells stably expressing SAM components (MCF10A-SAM cells), dCas9-VP64 and MCP-p65-HSF1 (gift from Feng Zhang, Addgene #61425 and #61426), were infected with the lentiviral SAM sgRNA library with a low titer of MOI=0.5. A library coverage of at least 500-fold was maintained throughout the experiment.

To screen for suppressors of YAP5SA activity, the MCF10A-SAM cells expressing the SAM library were superinfected with YAP5SA and cultured for two weeks to allow outgrowth of cells expressing suppressors of YAP5SA. gDNA was isolated using the QIAamp DNA Blood Maxi Kit (Qiagen). Sequencing libraries were generated using nested PCR. In the first PCR reaction, the integrated sgRNA cassette was amplified and then the second PCR was performed to add Illumina adapters and barcodes for NGS (primer sequences are listed in Supplementary Table 6). Libraries were quantified with the Agilent 2100 Bioanalyzer automated electrophoresis system (Agilent Technologies) and subjected to 75 bp single-end Illumina Sequencing on a NextSeq500. Reads were extracted in FastQ format using bcl2fastq v1.8.4 (Illumina).

### SAM analysis

Quality-filtering (>Q30) of the sequencing data and adapter removal was performed with Cutadapt (v2.7). The filtered reads were mapped to a custom reference containing all sgRNA sequences in the SAM library using Bowtie2. To all samples a pseudocount of +1 was added to avoid division by 0. The reads were normalized by sequencing depth (reads per sgRNA/ million mapped reads) and subsequently used in RSA analysis. For the RSA analysis, the following parameters were used, based on the median and the standard deviations (SDs) of the reads: --l = median plus 1xSD, --u= median plus 3×SD.

### Western blotting

Cell lysates were prepared using RIPA buffer (50 mM Hepes pH 7.9, 140 mM NaCl, 1 mM EDTA, 1% Triton X-100, 0.1% Na-deoxycholate, 0.1% SDS) complemented with sodium pyrophosphate and protease inhibitor cocktail (Sigma). Lysates were cleared by centrifugation and denatured in electrophoresis sample buffer at 95°C for 5 min. Proteins were separated on 8% Bis-Tris gels and transferred onto a 0.45 µm PVDF membrane (Millipore). Membranes were blocked with 5% skim milk powder in TBS, probed with primary antibodies diluted in 5% BSA in TBS-T and subsequently incubated with the appropriate horseradish peroxidase-coupled secondary antibodies. Visualization was performed using chemiluminescence HRP substrate (Clarity Western ECL Substrate, Bio-Rad).

### Crystal violet staining

For crystal violet staining, MCF10A cells were grown in triplicates on 6-well dishes, fixed with 3.7% paraformaldehyde for 10 min, stained with 0.1% crystal violet (Sigma-Aldrich) in 20% ethanol for 30 min and photographed.

### Co-immunoprecipitation

For exogenous co-immunoprecipitation to analyse homo- and heterodimerization of JUN, 293T cells were transfected using PEI-Max (Sigma-Aldrich). 48 h after transfection, cells were lysed in RIPA buffer containing protease inhibitor cocktail (Sigma-Aldrich). Immunoprecipitation of Flag-tagged proteins from cleared lysates was performed with Anti-Flag M2 Affinity Agarose Gel (Sigma-Aldrich) at 4°C for 3 h. Immunoprecipitates were washed three times with RIPA buffer and Flag-tagged proteins were eluted by two consecutive elution steps with 400 µg/ml Flag-peptide (Sigma-Aldrich) for 30 min at 4°C. Eluates were boiled with sample buffer and subjected to immunoblotting.

### RNA-Sequencing

RNA-Sequencing was performed as described previously (Kim et al., Nat. Commun., 2022). Briefly, total RNA was extracted using RNeasy® Micro Kit (Qiagen) with on-column DNaseI (Qiagen) digestion. RNA integrity (all processed samples had a RIN>8) was verified with the Agilent Bioanalyzer 2100 automated electrophoresis system (Agilent Technologies). mRNA was isolated using the NEBNext® Poly(A) mRNA Magnetic Isolation Module (NEB) from 1 µg of total RNA and library preparation was conducted with the NEBNext® Ultra RNA Library Prep Kit for Illumina (NEB) with Dual Index Primers (NEBNext® Multiplex Oligos for Illumina, NEB) following the manufacturer’s description. Cycles for amplification of the cDNA were determined by qRT-PCR. Libraries were quantified with the Agilent 2100 Bioanalyzer automated electrophoresis system (Agilent Technologies) and subjected to 75 bp single-end Illumina Sequencing on a NextSeq500. Reads were extracted in FastQ format using bcl2fastq v1.8.4 (Illumina). For all RNA-Sequencing samples, three biological replicates per condition were analyzed.

### RNA-Sequencing analysis

Adapter removal, size selection (reads > 25 nt) and quality filtering (Phred score > 43) of FASTQ files were performed with Cutadapt. Reads were then aligned to human genome (hg19) using Bowtie2 (v2.2.9) with default settings. Read count extraction was performed in R using countOverlaps (GenomicRanges). Differential gene expression analysis was done with DESeq2 (v3.26.8) using default parameters. Cluster analysis of YAP target genes was performed by MFuzz in R using three clusters.

### Statistics and reproducibility

All statistical analyses were performed in R (v4.1.0). The graphs always display the mean value and the standard error of the mean (SEM) unless stated otherwise. The statistical test performed is always given in the respective Figure legend.

### qRT-PCR

Total RNA was extracted with peqGOLD TriFast Reagent (VWR). First-strand cDNA synthesis was performed from 1 µg RNA using M-MLV Reverse Transcriptase (Promega) and random hexamer primers (Sigma-Aldrich) following the manufactureŕs instructions. qPCR reactions were conducted in technical triplicates using InnuMIX qPCR DSGreen Standard Mix (Analytik Jena) on a StepOnePlus™ Real-Time PCR System (Thermo Fisher Scientific). Expression values were normalized to *B2M* as housekeeping gene using the ddCt method. Primer sequences are listed in Supplementary Table 6.

### SLAM-Seq

For SLAM-Seq, MCF10A JUN knockout cells expressing JUN-AID-V5 fusion protein and Transport inhibitor response 1 (TIR1) were grown to 60-70% confluency and treated with 100 µM indole-3-acetic acid (IAA, Sigma-Aldrich) for 1h to deplete JUN protein. Newly synthesized RNA was then labelled by incubating the cells for 1h with 100 µM 4-thiouridine (4-sU, Sigma-Aldrich). RNA extraction was performed with peqGOLD TriFast Reagent (VWR) and total RNA was alkylated for 15 min using 10 mM iodoacetamide (Sigma-Aldrich). 3’-end mRNA sequencing libraries were generated from 335 ng alkylated RNA using the QuantSeq 3′ mRNA-Seq Library Prep Kit for Illumina (Lexogen). 75bp single-end sequencing was performed on a NextSeq500 Illumina sequencer.

### SLAM-Seq analysis

SLAM-Seq was analysed by a Nextflow SLAM-Seq pipeline (https://nf-co.re/slamseq). In parallel, a standard RNA-Seq analysis was performed to infer gene expression changes of the steady-state pool. Log2-fold changes of the SLAM-Seq data (based on T-to-C conversions) and the steady-state pool upon auxin-dependent JUN-AID degradation were inferred by DESeq2.

### Gene set enrichment analysis

Gene set enrichment was performed using the MSigDB GSEA tool (v 4.3.1) with a GSEAPreranked analysis.

### Hydrodynamic tail vein injection

Hepatocellular carcinoma was induced in wildtype C57BL/6J mice by delivering Sleeping Beauty (SB) transposon system into the livers of 6 weeks old male mice via hydrodynamic tail vein injection (HDTVI). Injection cocktails contained 10 µg pCMV(CAT)T7-SB100 (gift from Zsuzsanna Izsvak; Addgene # 34879); 25 µg pSBbi-Myr-Akt-HA and 25 µg pSBbi-eGFP-IRES-V5-YAP5SA/ pSBbi-cJUN-WT-IRES-V5-YAP5SA/ pSBbi-cJUN-M14-IRES-V5-YAP5SA.

All plasmids were purified using the Endotoxin-free plasmid DNA purification kit (Macherey-Nagel), the total volume was adjusted to 10% (ml) of the body weight (grams) using sterile Ringeŕs lactate solution (WDT) and injected into the lateral tail veins of the mice.

### Histology and immunohistochemistry of mouse liver cancer sections

For histological analyses, formalin-fixed paraffin-embedded mouse livers were sectioned at 5 μm and stained with hematoxylin and eosin (H&E). For immunohistochemical analyses, liver sections were stained with antibodies against HA-tag, YAP and JUN (Supplementary Table 6) following the standard procedures. In brief, after deparaffinization and rehydration, antigen retrieval was done by boiling the slides in citrate buffer pH 6.0 (Abcam). Endogenous peroxygenase was blocked using 3% (v/v) H_2_O_2_ in PBS. After blocking with 5% (w/v) BSA in PBS-T, slides were incubated with primary antibodies in a humidified container at 4 °C overnight. Incubation with appropriate horseradish peroxidase-coupled secondary antibodies was performed at RT for 2h. Visualization was done using Vector® ImmPACT DAB Peroxidase Substrate (Vector Laboratories), followed by a hematoxylin counterstaining. The slides were imaged using a slide scanner Axio Scan.Z1 microscope (Zeiss).

### Quantification of tumor load

To quantify tumor load from H&E stained sections of mouse livers, the area of all tumor nodules per liver was measured (in pixels) and divided by the total liver area. Quantification was performed using FIJI. The analysis was performed in a blinded manner and unblinded after the analysis was complete.

### BioID affinity purification and preparation for MS

MCF10A JUN knockout cells stably expressing BirA*-c-JUN fusion proteins were treated with 50 μM biotin (Sigma-Aldrich) for 18 h; 1.5 × 10^7^ cells were collected per sample, snap frozen in liquid nitrogen and stored at −80 °C till further use. Each cell pellet was resuspended in 4.75 ml lysis buffer (50 mM Tris pH 7.5, 150 mM NaCl, 1 mM EDTA, 1 mM EGTA, 1% Triton-X100, 0.1% SDS, 1mg/ml aprotinin, 0.5 mg/ml leupeptin, 250 U turbonuclease) and rotated for 1 h at 4°C. The samples were then sonicated (Bioruptor Plus, Diagenode) for 5 cycles (60 sec ON/30 sec OFF) at high setting and 20°C. Cell debris was removed by centrifugation (30 min at 4°C and 17.000×g). Streptavidin Sepharose High Performance Beads (Cytiva) were acetylated by adding 20 mM sulpho-NHS acetate twice for 30 min at RT. The reaction was quenched using 1 M Tris pH 7.5 and the beads were washed extensively with PBS. The acetylated streptavidin beads were equilibrated in lysis buffer, added to the lysate and incubated at 4°C for 3 h with rotation. After extensive washing with 40 mM ammonium bicarbonate, samples were digested with 1 µg LysC overnight at 37°C. Peptides were eluted with 150 µl 50 mM ammonium bicarbonate twice and digested with 1 µg trypsin. For elution of the biotinylated peptides, the beads were briefly mixed twice with 150 µl of 80% ACN and 20% TFA. Eluates were dried, resuspended in 200 mM HEPES pH 7.5 and trypsin (1 µg) was added to digest the peptides. Desalting and purification were performed using Waters Oasis® HLB µElution Plate 30 µm (Waters Corporation) according to the manufacturer’s instructions. Briefly, the columns were conditioned with 3×100 µl OASIS Buffer B (80% (v/v) acetonitrile; 0.05% (v/v) formic acid) and equilibrated with 3×100 µl OASIS Buffer A (0.05% (v/v) formic acid in Milli-Q water). The samples were loaded on the column, washed three times with 100 µl solvent A and then eluted with 50 µl OASIS buffer B twice. Eluates were dried using a speed vacuum centrifuge, resuspended in MS buffer A (5% acetonitrile, 0.1% formic acid) and loaded onto Evotips (Evosep) according to the manufacturer’s instructions. Briefly, the Evotips were washed with Evosep buffer B (0.1% formic acid in acetonitrile), conditioned with 100% isopropanol and equilibrated with Evosep buffer A (0.1% acetonitrile). Subsequently, the samples were loaded onto the Evotips and washed with Evosep buffer A. The loaded Evotips were filled up with buffer A and stored until measurement.

### MS analysis

Peptides were separated using the Evosep One system (Evosep) equipped with an 8 cm x 150 μm i.d. packed with 1.5 μm Reprosil-Pur C18 beads column (Evosep Endurance, EV-1106, PepSep). Samples were run with a pre-programmed proprietary Evosep gradient of 21 min (60 samples per day, 60SPD) with water, 0.1% formic acid, solvent B acetonitrile and 0.1% formic acid as solvents. The LC was coupled to an Orbitrap Exploris 480 (Thermo Fisher Scientific) using PepSep Sprayers and Proxeon nanospray source. The peptides were introduced into the mass spectrometer via a PepSep Emitter 360-μm outer diameter × 20-μm inner diameter, heated to 300 °C, and a spray voltage of 2.2 kV was applied. The temperature of the injection capillary was set to 300°C and the radio frequency ion funnel to 30%. For DIA data collection, full scan mass spectrometric (MS) spectra with mass range 350-1650 m/z were collected in profile mode in Orbitrap with a resolution of 120,000 FWHM. The default charge state was set to 2+. The fill time was set to a maximum of 45 ms with a limitation of 3 × 10^6^ ions. DIA scans were recorded with 35 mass window segments of different widths across the MS1 mass range. Higher collisional dissociation fragmentation (stepped normalised collision energy; 25, 27.5 and 30%) was applied and MS/MS spectra were acquired at a resolution of 15,000 FWHM with a fixed first mass of 200 m/z after accumulation of 1 × 106 ions or after 37 ms filling time (whichever occurred first). Data was collected in profile mode and processed using Xcalibur 4.5 (Thermo Fisher Scientific) and Tune version 4.0.

### Analysis of BioID data

Raw DIA data were analysed using the directDIA pipeline in Spectronaut (v.16, Biognosysis) with BGS settings, except the following parameters: Imputation strategy = Global Imputing, Protein LFQ method = QUANT 2.0, Proteotypicity Filter = Only protein group specific, Major Group Quantity = Median peptide quantity, Minor Group Quantity = Median precursor quantity, Data Filtering = Q-value percentile (0.2), Normalisation strategy = Global Normalisation on median, Row Selection = Identified in All Runs. Data were searched using a species-specific (*Homo sapiens*, 20.186 entries) and a contaminant database (247 entries) from Swissprot. Data were searched with the following variable modifications: Oxidation (M), Acetyl (Protein N-term) and Biotin (K). A maximum of 2 missed cleavages for trypsin and 5 variable modifications were allowed. Identifications were filtered to achieve a 1% FDR at the peptide and protein levels. Relative quantification was performed in Spectronaut for each paired comparison using the replicate samples from each condition. The data (candidate table) and data reports (protein quantities) were then exported, and further data analyses and visualisation was performed with Rstudio using in-house pipelines and scripts. A log2FC cutoff of 0.58 and a q-value <0.05 were set to select significant proteins.

### CUT&RUN

CUT and RUN experiments were performed as described previously (Kim et al., Nat. Commun., 2022). Briefly, for each CUT and RUN reaction 200,000 cells were trypsinized, washed, resuspended in 100 µl wash buffer (20 mM HEPES, pH7.5, 150 mM NaCl, 0.5 mM Spermidine) and bound to 10 µl activated BioMag®Plus Concanavalin A magnetic beads (Polysciences) for 10 min at room temperature. The cells were then incubated with antibodies diluted in 100 µl antibody buffer (wash buffer + 0.01% digitonine and 2 mM EDTA) overnight at 4°C. IgG rabbit antibody was used as negative control. After incubation with antibodies, the beads were washed in digitonin wash buffer (wash buffer + 0.01% digitonin) and incubated 1 h at 4°C with 1 µg/ml protein A/G Micrococcal Nuclease fusion protein (pA/G MNase). After washing with digitonin wash buffer, beads were rinsed with low salt buffer (20 mM HEPES, pH7.5, 0.5 mM Spermidine, 0.01% digitonine), resuspended in 200 µl incubation buffer (20 mM HEPES, pH7.5, 10 mM CaCl2, 0.01% digitonin) and placed at 0°C to initiate cleavage. After 30 min, reactions were stopped by adding 200 µl STOP buffer (170 mM NaCl, 20 mM EGTA, 0.01% digitonin, 50 µg/ml RNAse A) and the samples were incubated 30 min at 37°C to digest the RNA and release the DNA fragments.

The samples were then treated with proteinase K for 1 h at 50°C and the DNA was purified using Phenol/Chloroform/Isoamyl alcohol. After precipitation with glycogen and Ethanol, the DNA pellet was resuspended in 0.1 X TE and used for DNA library generation with the NEBNext^®^ Ultra™ II DNA Library Prep Kit for Illumina^®^ (New England Biolabs). Adaptor ligation was performed with 1:25 diluted adaptor and 15 cycles were used for library amplification using dual indices (NEB dual index kit). Paired-end 2×25 bp sequencing was performed on a NextSeq500 Illumina Sequencer.

### CUT&RUN analysis

Adapter removal and quality trimming was performed by Cutadapt. Since the carry-over DNA of pAG-MNase purification from *E. coli* was used as spike-in control, mapping was performed to hg19 and a human repeat-masked *E. coli* genome by Bowtie2. Paired-end reads mapped to hg19 with inserts <120 bp were extracted using alignment Sieve (deepTools). A scaling factor was inferred by: scaling factor= ^mapped reads < 120 bp to hg19^/_mapped reads to *E. coli*_

The scaling factor was used to generate spike-in normalized bigWig files by bamCoverage (deepTools). For peak calling, SEACR was used in the “stringent” mode and the peaks of the individual replicates were intersected by ChIPpeakAnno in R. Only peaks occurring in all replicates were retained for further analysis to generate a conservative peak set. For quantitative analyses, the spike-normalized bigWig files were used in computeMatrix reference-point (DeepTools), e.g. using peaks as reference point. The matrix output of computeMatrix, was then used for further analyses in R. Heatmaps were generated by plotHeatmap (DeepTools) using the output of computeMatrix.

### Data Availability

The Next-generation sequencing data generated in this study have been deposited in the GEO database under accession code GSE235968 https://www.ncbi.nlm.nih.gov/geo/query/acc.cgi?acc=GSE235968

## Acknowledgements

B.v.E. was supported by grants from the BMBF (16GW0271K), DFG (EY 120/4-1), and the German Cancer Aid (Deutsche Krebshilfe; 70113138). The FLI is a member of the Leibniz Association and is financially supported by the Federal Government of Germany and the State of Thuringia. The DNA Sequencing, the Proteomics, and the Flow Cytometry core facilities as well as the core service histology of the FLI are gratefully acknowledged. We would like to thank all the members of the von Eyss lab, von Maltzahn lab, and Kaether lab for helpful discussion and Christin Ritter and Tom Hünniger and all the animal care takers at FLI for excellent technical support. Some of the figures were created with BioRender.com

## Author Contributions

Conceptualization, Y.K., A.C.V., M.J., B.v.E.; Methodology, Y.K., A.C.V., M.J., B.v.E.; Software, B.v.E.; Formal analysis, Y.K., A.C.V., M.J., B.v.E.; Investigation, Y.K., A.C.V., M.J., B.v.E.; Data Curation, B.v.E.; Writing – Original Draft, A.C.V., M.J., B.v.E.; Visualization, Y.K., A.C.V., M.J., B.v.E.; Supervision, B.v.E..

